# Hepatocyte-specific activity of TSC22D4 triggers progressive NAFLD by impairing mitochondrial function

**DOI:** 10.1101/2021.12.17.473222

**Authors:** Gretchen Wolff, Minako Sakurai, Amit Mhamane, Adriano Maida, Ioannis K. Deligiannis, Kelvin Yin, Peter Weber, Annika Weider, Maria Troullinaki, Anja Zeigerer, Michael Roden, Nadine Volk, Tanja Poth, Thilo Hackert, Lena Wiedmann, Francesca De Angelis Rigotti, Juan Rodriguez-Vita, Andreas Fischer, Rajesh Mukthavaram, Pattraranee Limphong, Kiyoshi Tachikawa, Priya Karmali, Joseph Payne, Padmanabh Chivukula, Bilgen Ekim-Üstünel, Celia P. Martinez-Jimenez, Julia Szendrödi, Peter Nawroth, Stephan Herzig

## Abstract

**Objective:** Fibrotic organ responses have recently been identified as long-term complication in diabetes. Indeed, insulin resistance and aberrant hepatic lipid accumulation represent driving features of progressive non-alcoholic fatty liver disease (NAFLD), ranging from simple steatosis and non-alcoholic steatohepatitis (NASH) to fibrosis. Effective pharmacological regimens to stop progressive liver disease are still lacking to-date.

**Methods:** Based on our previous discovery of transforming growth factor beta-like stimulated clone (TSC)22D4 as a key driver of insulin resistance and glucose intolerance in obesity and type 2 diabetes, we generated a TSC22D4-hepatocyte specific knockout line (TSC22D4-HepaKO) and exposed mice to control or NASH diet models. Mechanistic insights were generated by metabolic phenotyping and single cell liver sequencing.

**Results:** Hepatic TSC22D4 expression was significantly correlated with markers of liver disease progression and fibrosis in both murine and human livers. Indeed, hepatic TSC22D4 levels were elevated in human NASH patients as well as in several murine NASH models. Specific genetic deletion of TSC22D4 in hepatocytes led to reduced liver lipid accumulation, improvements in steatosis and inflammation scores and decreased apoptosis in mice. Single cell RNA sequencing revealed a distinct gene signature identifying an upregulation of mitochondrial-related processes. An enrichment of genes involved in the TCA cycle, mitochondrial organization, and triglyceride metabolism underscored the hepatocyte-protective phenotype and overall decreased liver damage as seen in mouse models.

**Conclusions:** Together, our data uncover a new connection between targeted depletion of TSC22D4 and intrinsic metabolic processes in progressive liver disease. Cell-specific reduction of TSC22D4 improves hepatic steatosis, inflammation and promotes hepatocyte survival thus paving the way for further preclinical therapy developments.

## INTRODUCTION

With more than 1.5 billion overweight subjects worldwide, the prevalence of obesity-related metabolic complications is steadily increasing and imposes continuous pressure on global health care systems. In this regard, non-alcoholic fatty liver disease (NAFLD) is commonly regarded as the hepatic manifestation of the obesity-driven metabolic syndrome. Indeed, fatty liver disease characterizes more than 60 % of obese patients, and NAFLD has recently been identified as a main predictor for diabetes risk in humans with prediabetes [1]. Importantly, progression of steatosis to non-alcoholic steatohepatitis (NASH) and eventually fibrosis, represents the key etiology for the development of liver cancer, starting to outpace today’s classical, virus-related causes for liver cancer [2]. Despite its clinical importance, effective therapeutic modalities against NAFLD have not been established to date. This can at least partially be explained by the involvement of different hepatic cell types in its pathogenesis and the resulting signaling complexity. In fact, progressive liver disease results from the intricate interplay between distinct hepatic cell populations, including inflammatory cells, hepatic stellate cells and hepatocytes [3]. Indeed, a number of secreted mediators that drive the mutual changes in cellular phenotypes upon NASH progression have been identified, e.g., TGFβ [2; 4]. Whether common regulatory nodes exist in different hepatic cell populations that may contribute to progressive liver dysfunction remains unknown.

Transforming growth factor stimulated clone (TSC)22D4 is a globally expressed protein and belongs to the TSC protein family, which share a TSC box containing a leucine zipper motif mediating homodimerization or heterodimerization with other family members [5]. Members of the TSC family play diverse roles such as in glucocorticoid and stress signaling, cell proliferation, and apoptosis, yet the exact mechanism of action remains unknown [6-9]. However, we have recently shown that livers from mice with cancer cachexia have elevated TSC22D4 expression levels, positively correlating with the degree of tissue wasting and dysregulated lipid metabolism [10]. Follow-up studies demonstrated elevated hepatic levels of TSC22D4 also in obese patients with type 2 diabetes, particularly linked to insulin resistance [11]. Indeed, initial proof-of-principle studies highlighted the utility of AAV-driven continuous inhibition of TSC22D4 to counteract diabetes-related metabolic phenotypes in mouse models of type 2 diabetes. Hepatic knockdown of TSC22D4 not only ameliorated insulin resistance but also improved blood glucose levels without any adverse changes in body mass [11]. Overall, these studies positioned hepatic TSC22D4 as a potentially druggable upstream regulator of both lipid and glucose metabolism in type 2 diabetes. However, whether TSC22D4 beyond its acute impact on key metabolic parameters may also exert control in long-term diabetic complications, particularly liver fibrosis, remained unclear but would underscore its pharmacological attractiveness.

Here, we show that TSC22D4 correlated with fibrosis in both murine and human liver samples. Depletion of TSC22D4 in hepatocytes improved liver lipid accumulation in mice fed NASH-promoting diets and reduced liver injury. Furthermore, single cell RNA sequencing of hepatocytes revealed genes involved in enhanced mitochondrial efficiency, lipid metabolism, oxidative phosphorylation under conditions of progressive fatty liver disease.

## MATERIALS AND METHODS

### ANIMALS

Mice were housed at room temperature with 12-h light-dark cycle on control diet (Research diets, New Brunswick, NJ, USA, D12450B), 60% high fat diet (Research diets, D12492i), control for MCD (Research diets, A02082003BY), MCD (Research diets, A02082002BR) or chow for weeks as indicated in particular study. Mice had *ad libitum* access to food and water and were weighed weekly and inspected daily for general health. At the conclusion of each study, tissues were flash frozen for analysis or placed in histofix (4% formaldehyde, Carl Roth GmbH, Karlsruhe, Germany) for immunohistochemistry. All experiments were performed in accordance with the European Union directives and the German animal welfare act (Tierschutzgesetz) and were approved by local authorities (Regierungspräsidium Karlsruhe, license #G117/18 or 18 or the Government of Upper Bavaria, license #ROB-55.2-2532.Vet_02-17-49). *Carbon tetrachloride (CCl*_*4*_,*) experiment*, 9-week-old, male Balb/c mice were purchased from Janvier Labs (Le Genest-Saint-Isle, France). CCl_4_ was diluted 1:5 in olive oil (Carl Roth 8873.1) and injected intraperitoneally (i.p.) at a concentration of 0.5 µl/g body weight, (VWR International SIAL289116) 3x weekly for two weeks. Tissues were collected as described above. STAM™ 10 week old liver tissues in figure 2 were purchased from SMC Laboratories, Inc. (Tokyo, Japan).

NASH and fibrosis studies were performed with Albumin-cre mice crossed to the TSC22D4 floxed/floxed line (C57Bl/6N background) (Supp Fig. 1). The targeting strategy was carried out by Taconic Biosciences GmbH (Cologne, Germany) based on NCBI transcript NM_023910.6. A targeting vector containing genomic fragments of the mouse TSC22D4 locus as well as the loxP flanked region was generated. LoxP sites flanked the genomic region containing exons 1 and 2 and 1.5 kb upstream of exon 1 (promoter region).

#### Genotyping

The presence of the floxed allele was confirmed by PCR performed using DNA extracted from mouse earclip biopsies. Expected PCR products were 266bp for the Tsc22d4 wild-type allele and 385 for the floxed allele. For routine genotyping, primers flanking the loxP site located to the right of exon 2 were used (forward CTTCCCTGATTTCAGGTGATG and reverse ATCCCACACTCCAGGAGAAG). For Albumin-cre crossed mice, a PCR 320bp fragment identifying the presence of cre was observed (forward primer GAACCTGATGGACATGTTCAGG and reverse primer AGTGCGTTCGAACGCTAGAGCCTGT).

#### STAM experiments

Male mice with hepatocyte-specific deletion of TSC22D4 or cre^-^ littermates were treated as previously published [12]. Briefly, 2-day old pups were injected with a low dose of streptozotocin (Sigma) 200 µg i.p., randomized at 3 weeks of age and placed on control (CD) or high fat diet (HFD). Mice continued on diet for 12 weeks until tissue collection as described above.

*For MCD diet studies*, male, hepatocyte-specific TSC22D4 Ko mice and Wt littermates at 10 weeks of age were fed MCD or matched MCD control diet for 3 weeks and tissues collected as described above.

### LIPID NANOPARTICLE FORMULATION

#### Preparation of LUNAR-siRNA Nanoparticles

TSC22D4 siRNA (sense: 5’-UNA-G/mGrAmCrGmUrGmUrGrUrGmGrAmUrGmUrUmUrA/UNA-U/mU-3’; antisense: 5’-mUrAmArAmCrAmUrCmCrAmCmAmCrAmCrGmUrCmC/UNA-U/mU-3’, wherein rN is RNA, mN is 2’-O-methyl RNA, and UNA is unlocked nucleic acid), and non-targeting control siRNA (sense: 5’-UNA-U/mArGmCrGmArCmUrArArAmCrAmCrAmUrCmGrC/UNA-U/mU-3’, antisense: 5’-UNA-G/rCmGrAmUrGmUrGmUrUmUmAmGrUmCrGmCrUmA/UNA-U/mU-3’, wherein rN is RNA, mN is 2’-O-methyl RNA, and UNA is unlocked nucleic acid) were synthesized by Integrated DNA Technologies (Coralville, IA). ATX lipid was synthesized as described previously [13]. LNPs were prepared as described previously [14] by mixing appropriate volumes of lipids in ethanol with an aqueous phase containing siRNA duplexes using a Nanoassemblr microfluidic device, followed by downstream processing. For the encapsulation of siRNA, desired amount of siRNA (TSC22D4 or non-targeting control siRNA) was dissolved in 5 mM citrate buffer (pH 3.5). Lipids at a molar ratio of 58% ionizable ATX lipid, 7% DSPC (1,2-distearoyl-sn-glycero-3-phosphocholine) (Avanti Polar Lipids), 33.5% cholesterol (Avanti Polar Lipids), and 1.5% DMG-PEG (1,2-dimyristoyl-sn-glycerol, methoxypolyethylene glycol; PEG chain molecular weight: 2000) (NOF America Corporation) were dissolved in ethanol. At a flow ratio of 1:3 ethanol/aqueous phases, the solutions were combined in the microfluidic device using Nanoassemblr (Precision NanoSystems). The total combined flow rate was 12 mL/min per microfluidics chip and the total lipid-to-RNA weight ratio was ∼30:1. The mixed material was then diluted three times with Tris buffer (pH 7.4) containing 50 mM sodium chloride and 9% sucrose after leaving the micromixer outlet, reducing the ethanol content to 6.25%. The diluted LNP formulation was concentrated and diafiltered by tangential flow filtration using spectrum hollow fiber membranes (mPES Kros membranes, Repligen) and Tris buffer (pH 7.4) containing 50 mM sodium chloride and 9% sucrose. A total of 10 diavolumes were exchanged, effectively removing the ethanol. The particle size and polydispersity index (PDI) were characterized using a Zen3600 (Malvern Instruments, with Zetasizer 7.1 software, Malvern, U.K.). Encapsulation efficiency was calculated by determining the unencapsulated siRNA content by measuring the fluorescence intensity (Fi) upon the addition of RiboGreen (Molecular Probes) to the LNP and comparing this value to the total fluorescence intensity (Ft) of the RNA content that is obtained upon lysis of the LNPs by 1% Triton X-100, where % encapsulation = (Ft − Fi)/Ft × 100). Encapsulation efficiencies of these formulations were >90%.

### GENE EXPRESSION

RNA was prepared using the RNeasy micro kit (QIAGEN) including DNase treatment following the manufacturer’s protocol. Complementary DNA (cDNA) synthesis was performed with 200**–**1,000 ng total RNA using QuantiTect_ Reverse Transcription Kit (Qiagen). Quantitative real time polymerase chain reaction (qRT**–**PCR) was performed using TaqMan_ Gene Expression Assays (all Life Technologies, Darmstadt, Germany). TaqMan_Gene Expression Master Mix and the StepOnePlusTM Real-Time PCR System (Life Technologies) for TSC22D4 (Probe Mm00470231_m1). Normalization to Tbp/TBP (TATA box binding protein, Probe Mm01277042_m1) and calculation of relative expression values were performed with the ΔΔCT method.

### LIPID MEASUREMENTS IN MICE

Triglycerides (TR0100, Sigma Aldrich) and total cholesterol (CH200, Randox, Crumlin, UK) were measured in serum and liver lipid lysates using the Mithras Microplate Reader (Berthold Technologies GmbH & Co, Bad Wildbad, Germany) at 550 nm. Liver lipids were extracted at described previously [15].

### QUANTITATIVE NASH ASSESSMENT IN MICE

Liver samples were fixed with 4% Histofix (Carl Roth) at room temperature for 24 h then embedded in paraffin wax. Blocks were cut into sections 4 μm thick sections each and placed on glass slides. Sections were de-paraffinized with xylene and ethanol dilutions to rehydrate. Following rehydration and rinsing, slides were stained with Pikro-Sirius red (ref# 1422.00500, Morphisto GmbH, Vienna, Austria) for 60 min and dehydrated followed by mounting. For pathological assessment hematoxylin and eosin stained mouse liver sections (Leica ST5020 auto-stainer) were evaluated according to Kleiner DE, et al. Hepatology (2005), modified by Liang et al., PLOS (2014) and adopted for the present study consisting of a scoring strategy with features grouped into five broad categories: steatosis, inflammation, hepatocellular injury, fibrosis, and miscellaneous features [16; 17].

### HUMAN PATIENTS

#### Human NAFLD cohort

The prospective BARIA_DDZ study examines and follows obese individuals undergoing bariatric surgery and lean, healthy humans (controls) undergoing elective surgery such as cholecystectomy or herniotomy (registered clinical trial no. NCT01477957) [18; 19]. The cohort evaluated here consisted 33 females and 12 males with mean age of 42.5 years and mean BMI o f45.8 kg/m^2^. All participants were stratified based on histological assessment in groups without steatosis (NAFL−) or with steatosis (NAFL+) steatosis and inflammation (NASH) as described previously [18; 19]. All participants were examined using hyperinsulinaemic-euglycaemic clamps to determine peripheral and hepatic insulin sensitivity, and blood was sampled for routine lab parameters [18]. Participants maintained stable body weight for at least 2 weeks before the surgery. The study was approved by the Heinrich Heine University Düsseldorf Institutional Review Board and informed consent obtained in writing from all participants.

#### Human fibrosis cohort

Human liver samples were provided by the Tissue Bank of the National Center for Tumor Diseases (NCT) Heidelberg, Germany in accordance with the regulations of the tissue bank and the approval of the ethics committee of Heidelberg University [20]. The cohort evaluated here comprised 15 females and 24 males with and average age of 60.8 years and mean BMI 2 of 5.6 kg/m^2^. The human samples were obtained upon informed consent and study protocols adhered to the ethical guidelines of the updated version of the Declaration of Helsinki approved by institutional review committee.

### RNA SEQUENCING

#### Single nuclei isolation

All mouse liver tissues were flash frozen in liquid nitrogen upon collection and then pulverized on dry ice without thawing using a tissue lyser, (Qiagen). Briefly, pulverized liver was placed into a pre-chilled Dounce homogenizer (Lab Logistics, #9651632) with 1 ml of homogenization buffer (HB) [(250 mM sucrose, 25 mM KCl, 5 mM MgCl2, 10 mM Tris buffer, 1 μM DTT), 1 x protease inhibitor tablet (Sigma Aldrich Chemie), 0.4 U/μL RNaseIn (Thermo Fisher Scientific), 0.2 U/μL Superasin (Thermo Fisher Scientific), 0.1% Triton X-100 (v/v) and 10 µg/mL Hoechst 33342 (Thermo Fisher Scientific) in RNase-free water], as described by Krishnaswami et al [21]. After 5 mins incubation in HB, we performed 5 slow strokes with the loose pestle, followed by 10 strokes with the tight tolerance pestle. The suspension was passed through a 50 μm sterilized filter (CellTrics, Symtex, #04-004-2327) into a 2 mL pre-chilled Eppendorf pre-chilled and an additional 1 mL of cold HB was used to wash the Dounce homogenizer and the filter. The sample was then centrifuged for 8 mins at 1000 x g in a swinging bucket centrifuge. The pellet was resuspended in 250 μL of pre-chilled HB and a density gradient centrifugation clean-up was performed for 20 min, using Iodixanol gradient (Optiprep, D1556, Sigma Aldrich Chemie). The final pellet was resuspended gently in 200 µL of nuclei storage buffer (NSB) (166.5 mM sucrose, 5 mM MgCl2, 10 mM Tris buffer pH 8.0) containing additional RNase inhibitors 0.2 U/μL Superasin (Thermo Fisher Scientific, #AM2696) and 0.4 U/μL Recombinant RNase Inhibitor (Takara Clontech #2313A). The final suspension was filtered through a 35 μm cell strainer cap into a pre-chilled FACS tube prior to FACS sorting.

#### Flow cytometry

Prior sorting, 384-well PCR plates (thin-walled, BioRad, HSP3901) were freshly prepared with 810 nL of initial Lysis Buffer (LB1) using the Mosquito HV liquid handling robot (STP Labtech) as published previously [22]. Upon sorting, Hoechst stained diploid and tetraploid nuclei were singly sorted using a FACS sorter (BD FACSAria 3 sorter), 1 nucleus per well, with a 100 μm nozzle. Drop delay and cut-off point were optimized prior to every sorting, and the plate holder was maintained at 4 °C. Sorting accuracy was assessed using a colorimetric method with tetramethyl benzidine substrate (TMB, BioLegend, #421501) and Horseradish Peroxidase (HRP, Life Technologies, #31490) as described at Rodriguez et al [23]. The gating strategy used FSC-A/SSC-A to select intact nuclei stained with Hoechst, in which events above 10^4 are considered Hoechst positive. Subsequently, selection of singlets was performed by FSC-A/FSC-H followed by exclusion of doublets by FSC-A/FSC-W as previously described [24]. After sorting, every plate was firmly sealed (MicroAmp Thermo Seal lid, #AB0558), shortly vortexed for 10 s, centrifuged at 4°C, 2000 x g for 1 min, frozen on dry ice, and stored at −80°C until cDNA synthesis.

#### snRNA-seq2

The single-nucleus-RNA-seq2 methodology was used to capture a high number of transcripts from frozen tissues, allowing for the generation of double-stranded full-length cDNA as described by Richter *et al*. 2021. The SMART-Seq Single Cell Kit (SSsc; Cat. # 634472, Takara) was used and reaction volumes were miniaturized 6 times with the aid of the Mosquito HV robot as per the kit provider User Guide (cDNA synthesis using the mosquito HV genomics with the SMART-Seq® Single Cell Kit) addition of the second lysis buffer (LB2), containing NP40 2%, Triton-X100 1%, 1/400,000 diluted ERCC RNA spike-in, 3′ SMART-seq CDS Primer II A and RNAse-free water, results to a volume of 1.7 uL. Plates were immediately sealed, vortexed 20 s at 2000 rpm, centrifuged at 2000 x *g* for 30 s at 4ºC and incubated at 72 °C for 6 min. ERCC spike-ins (Thermo Fischer Scientific, Ref. 4456740) were diluted 1 in 10 with RNAse-free water, supplemented with 0.4 U/μL Recombinant RNase Inhibitor (Takara Clontech, Ref. 2313A). Fresh dilution of 1 in 400,000 was prepared immediately before the first strand synthesis. Reverse transcription and Pre-PCR amplification steps were followed as described by the manufacturer and the PCR program for the cDNA amplification was performed in a total of 20 cycles of: 1 min at 95 °C, [20 s at 95 °C, 4 min at 58 °C, 6 min at 68 °C] × 5, [20 s at 95 °C, 30 s at 64 °C, 6 min at 68 °C] × 9, [30 s at 95 °C, 30 s at 64 °C, 7 min at 68 °C] × 6, 10 min at 72 °C. We purified the amplified cDNA using Agencourt XP beads at 0,7X (Catalog No. A63881, Beckman) on the Mosquito HV, and then performed quantification of cDNA in an Agilent Bioanalyzer with a High Sensitivity DNA chip.

#### Library preparation

Sequencing libraries were prepared using the Illumina Nextera XT DNA Sample Preparation kit (Illumina, Ref. FC-131-1096) and the combination of 384 Unique Dual Indexes (Illumina-Set A to D, Ref. 20027213-20027216). Using the Mosquito HV robot, the reaction volumes of the Nextera XT chemistry were miniaturized by 10 times, following steps as previously described [25] In brief, 500 nL of undiluted cDNA (∼200pg/uL) were transferred to a new 384 well-plate containing 1500 nL of Tagmentation Mix (TD and ATM reagents). Accordingly, all Nextera XT reagents (NT, NPM and i5/i7 indexes) were added stepwise to a final library volume of 5 μL per well. The final PCR amplification was performed through 12 cycles. Once the libraries were prepared, 500 nL from each well were pooled together into a 1,5 mL tube to perform two consecutive clean-up steps with a ratio of sample to AMPure XP beads of 0,9X (Beckman Coulter, Ref. A63882). The final library sizes ranged between 200 and 1000 bp as assessed using a HS DNA kit in the Agilent Bioanalyzer. Prior to sequencing, the libraries were quantified using a Collibri library quantification kit (Thermo Fischer Scientific, Ref. A38524100) in a QuantStudio 6 Flex (Life Technologies).

#### Sequencing

Each of the pooled libraries from the 384 well plates were sequenced using an Illumina NovaSeq 6000 NGS sequencer in one lane of an SP300 Xp flowcell, in a paired-end 150 bases length resulting on an average of 1Million reads per nuclei.

### Hepatocyte isolation and transfection

Murine primary hepatocytes were isolated as described previously [26; 27]. Briefly, under anesthesia the visceral cavity was opened and the liver perfused via the venae cavae first with EGTA-containing HEPES/KH buffer followed by a collagenase-containing HEPES/KH buffer until liver digestion was visible. The liver was excised and placed in suspension buffer where hepatocytes were gently washed out. Cells were filtered through a 100-nm pore mesh, centrifuged and washed twice in suspension buffer. Hepatocytes were plated in collagen-coated 24-well plates (Thermo Fisher Scientific) in William’s Medium E (PAN-Biotech) containing 10% FCS (PAN-Biotech), 5% penicillin-streptomycin (Thermo Fisher Scientific) and 100 nM dexamethasone (Sigma-Aldrich) and maintained at 37 °C and 5% CO2. After 2 hours of attachment, cultures were washed with phosphate buffer saline (PBS) and incubated with 40nM LNP formulation with 2.4 µg/ml ApoE (Fitzgerald Acton, MA, U.S.A.) in William’s E medium for 6 hours, then washed with PBS and media was changed [26; 28].

### Seahorse assay

Hepatocytes were seeded at 30’000/well in 24-well seahorse plates that had been coated with a thin layer of collagen and maintained in William’s media E (PAN Biotech), 10% FBS (Sera Plus from PAN biotech), and 1% P/S, 100 nM Dexamethasone. 2 hours after seeing, cells were washed 2x with PBS then “transfected” with LNP-siRNAs with ApoE. LNPs (40nM) and ApoE (2.4ug/ml) were mixed in media (same one as above), kept at RT for 10min, then added to cells. 6 hours later media was replaced with fresh media and Seahorse assay performed 48 later. A mitostress test was performed according to the manufacturer’s instructions, with timed sequential injection of oligomycin 2uM, FCCP 1uM, and lastly rotenone+antimycin (1uM each). Oxygen consumption rate (OCR) data were normalised to cell number, as quantified using the CellQUANT cell proliferation Assay Kit (Invitrogen).

### STATISTICAL ANALYSIS

Data were graphed and analyzed using GraphPad Prism 9 software (La Jolla, CA, USA).

### STUDY APPROVAL

All experiments were performed in accordance with the European Union directives and the German animal welfare act (Tierschutzgesetz) and were approved by local authorities (Regierungspräsidium Karlsruhe, license #G117/18).

## RESULTS

### Hepatic TSC22D4 expression correlates with circulating lipids and markers of NASH and fibrosis in patients

To initially understand the relevance of TSC22D4 in persons with progressive liver disease, we investigated two cohorts. The first consisted of patients with steatosis or steatohepatitis (NASH), while the second had NASH and liver fibrosis. TSC22D4 mRNA levels were significantly increased in livers from obese persons both without (NAFL-) or with steatosis (NAFL+) and with histologically confirmed NASH (Fig. 1A). The levels of TSC22D4 positively correlated with serum triglycerides and negatively with HDL-cholesterol (Fig.1B) suggesting that increased hepatic levels of TSC22D4 associated with dyslipidemia in NAFLD. Additionally, consistent with our previous report [11], TSC22D4 mRNA positively correlated with increased glucose, insulin, and C-peptide measures (Supp. Fig. 2A-C). Patients with high levels of canonical fibrosis markers Col1α1, αSMA, and TGFβ, all had significantly elevated TSC22D4 mRNA expression, which correlated with fibrosis markers in these patients (Fig1. C-E). Together these data suggest a potential role for TSC22D4 in different stages of human NAFLD.

**FIG. 1.**
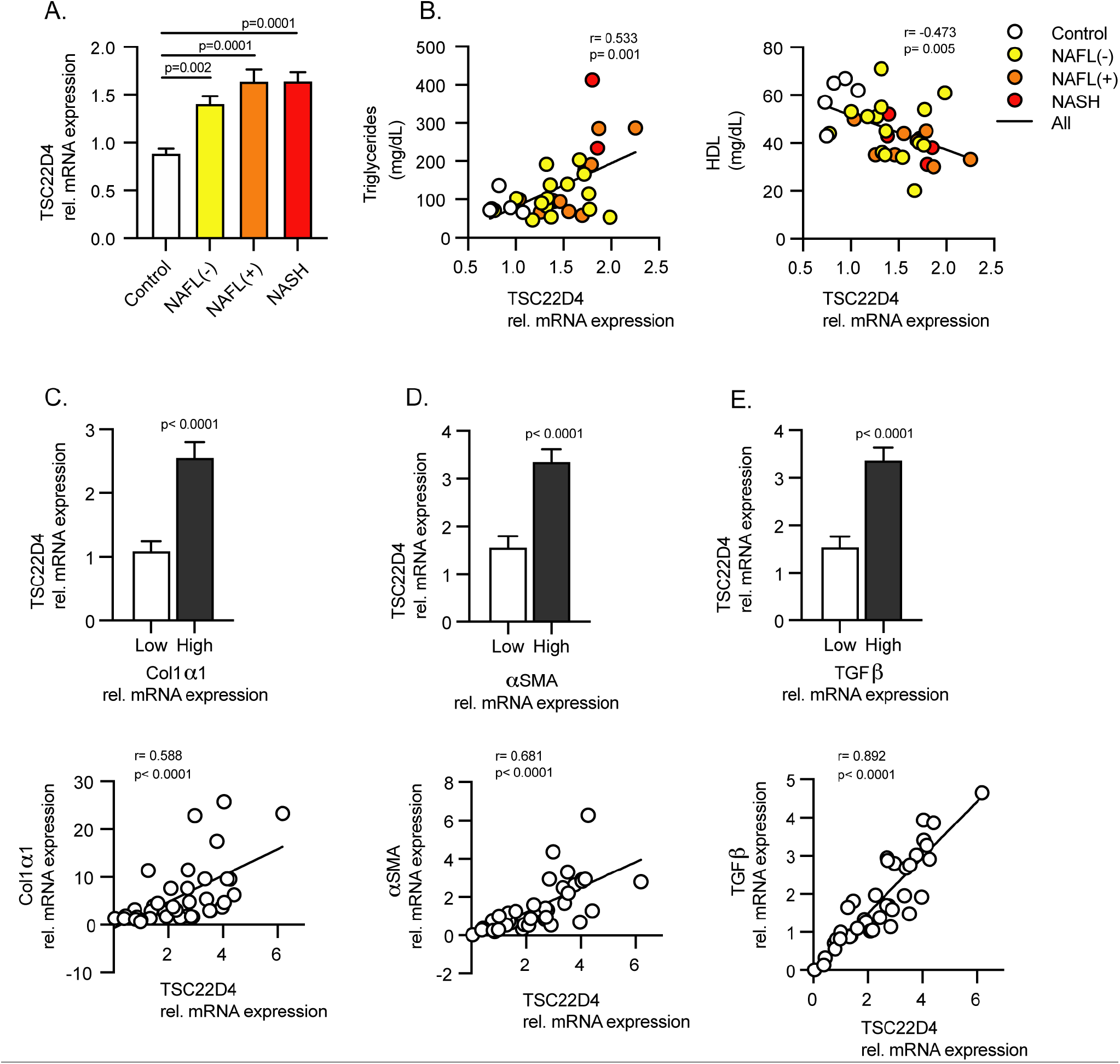
Hepatic TSC22D4 expression correlates with circulating lipids and markers of NASH and fibrosis in patients. Liver samples from patients classified as control, NAFLD negative, NAFLD positive, and NASH, (33 females, 12 males; mean age 42.5 years; mean BMI 45.8 kg/m^2^). TSC22D4 mRNA expression (A) and correlation with plasma triglycerides and HDL (B). NASH and fibrosis patient liver samples (15 females, 24 males; mean age 60.8 years; mean BMI 25.6 kg/m^2^) grouped based on expression of fibrosis markers, collagen 1α1, α-smooth muscle actin, or TGFβ. TSC22D4 mRNA levels in upper (high) and lower (low) 50% percentile of patients expressing and correlating with fibrotic markers collagen 1α1 (C) α-SMA (D) or TGFβ (E). Abbreviations: Nafl(-), patients without NAFLD diagnosis; Nafl(+), patients with NAFLD diagnosis; NASH, nonalcoholic steatohepatitis.

### TSC22D4 is upregulated in mouse models of fibrosis

CCl_4_ is a well-known potent chemical inducer of fibrosis in mice [29]. In agreement with results from patients presenting with clinical fibrosis, mice with a CCl_4_-induced fibrotic liver displayed increased TSC22D4 mRNA expression (Fig. 2A). Indeed, mice treated with CCl_4_ had a significant induction of several markers of liver fibrosis, TGFβ, αSMA and Col1α1 (Fig. 2B), and hepatic TGFβ significantly correlated with TSC22D4 mRNA levels (Fig. 2C). A more recent model used to recapitulate the NASH-fibrosis-hepatocellular carcinoma transition in mice is the STAM™ model [12]. STAM mouse livers also demonstrated a clear induction of TSC22D4 expression, which correlated with the induction of the pro-fibrotic marker TGFβ (Fig. 2D-E). Overall, these results support the hypothesis that the aberrant activation of TSC22D4 expression represents a common feature of both murine and human progressive liver disease.

**FIG. 2.**
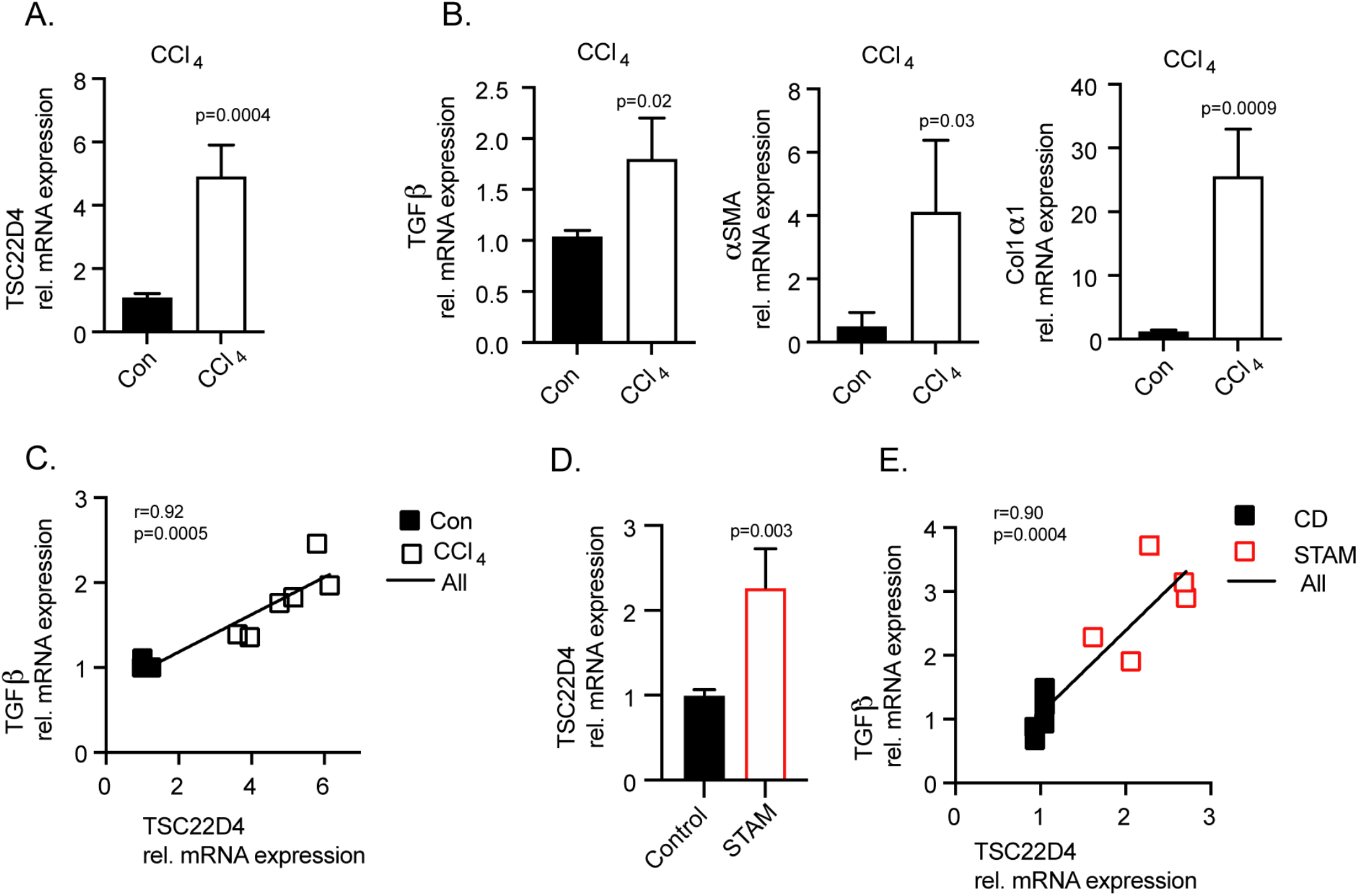
TSC22D4 is upregulated in mouse models of fibrosis. Carbon tetrachloride (CCl_4_) treated mice demonstrated significant upregulation of TSC22D4 mRNA (A) in conjunction with markers of fibrosis collagen 1α1, α-smooth muscle actin, or TGFβ (B). TSC22D4 correlated with marker of liver disease progression, TGFβ (C). STAM mice, streptozotocin-induced plus 9 w of HFD, exhibited increased TSC22D4 mRNA (D) that correlated with TGFβ expression (E) in liver. Abbreviations: HFD, high fat diet

### TSC22D4 knockout in hepatocytes decreases liver lipid accumulation in models of NASH and liver fibrosis in mice

Triglyceride accumulation within hepatocytes is a hallmark of NAFLD, and chronic steatosis leads to inflammation, apoptosis, and fibrosis [30]. The MCD diet model is a robust inducer of progressive NAFLD in mice [31]. In agreement with our earlier observation in human progressive NAFLD and STAM mice, animals fed the MCD diet had an increased expression of hepatic TSC22D4 (Fig 3A). When mice lacking TSC22D4 only in hepatocytes (Supp. Fig. 3A) were given the same diet, pathological examination revealed a decrease in liver lipid droplets and an improved sum of scores [16] relating to classic NASH characteristics (Fig. 3B-C). The improvement in liver injury was mainly the result of fewer apoptotic hepatocytes, reduced steatosis and inflammation scoring (Fig. 3D-E). In conjunction with the improved NASH phenotype seen in TSC22D4-HepaKO MCD fed mice we measured decreased levels of liver triglycerides and total cholesterol (Fig. 3F-G). TSC22D4-HepaKO mice exposed to the STAM model also showed decreased liver total cholesterol (Supp. Fig. 3B). TSC22D4 expression both positively correlated with the amount of liver triglycerides and negatively with serum HDL-cholesterol (Supp. Fig. 3C-D) as we observed in patient cohorts (Fig. 1B). Furthermore, TSC22D4 mRNA expression significantly correlated with TGFβ in both models of NAFLD (Supp. Fig. 3E-F) suggesting a robust connection between TSC22D4 and progressive liver disease. These experiments demonstrated that TSC22D4 contributed to key parameters of NAFLD progression, including dyslipidemia, inflammation, and hepatocyte apoptosis.

**FIG. 3.**
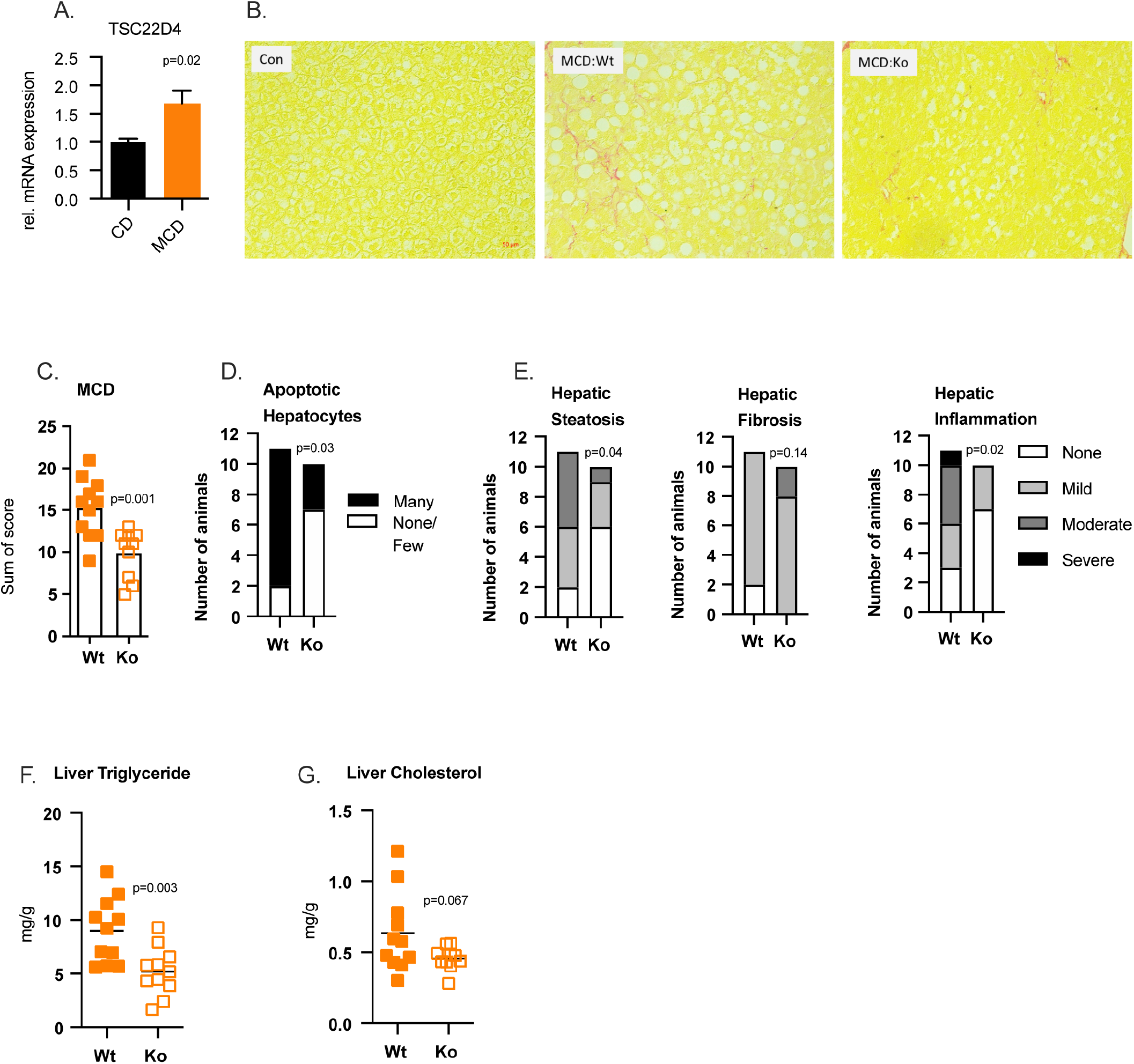
Hepatocyte specific loss of TSC22D4 decreases liver lipids in mice. Mice carrying TSC22D4 floxed allele with (Ko) or without (Wt) albumin cre-recombinase ensuring hepatocyte targeted deletion, were fed control or methionine-choline deficient diet for 3 weeks. Hepatic levels of TSC22D4 in Wt mice fed CD or MCD (A). Representative images of sirius red-stained liver tissue from CD and MCD fed mice (B). Hepatocyte-specific deletion of TSC22D4 reduced liver injury sum of scores (C) number of apoptotic hepatocytes (D) steatosis and inflammatory scores (E) on MCD diet. Liver total cholesterol (F) and triglycerides (G) were also reduced in Ko mice on MCD diet. Abbreviations: MCD, methionine-choline deficient; CD, control diet

### Loss of TSC22D4 in hepatocytes upregulates a mitochondrial-protective gene network in murine NAFLD

To gain mechanistic insight into the protective effect of hepatocyte-specific loss of TSC22D4 in NAFLD single cell RNA sequencing (scRNAseq) was performed on liver samples from TSC22D4-HepaKO and WT mice on MCD and control diet. T-distributed stochastic neighborhood embedding (tSNE) analysis revealed clustering of hepatocytes based on control or MCD diet whereas fewer cells were clustered within diet based on TSC22D4-HepaKO and WT (Fig. 4A) but with no batch effect (Supp. Fig. 4A). Subsequent gene ontology (GO) analysis identified gene signatures related to mitochondrial maintenance, the TCA-cyle and electron transport chain as well as cellular respiration, triglyceride metabolism, and lipid catabolic processes among the top 100 significantly enriched pathways (Fig. 4B). TSC22D4-HepaKO mice on control diet presented a slightly stronger clustering compared to WT and here the top 10 GO biological pathways also included metabolic and catabolic processes as well as DNA repair and autophagy (Supp. Fig. 4B).

**FIG. 4.**
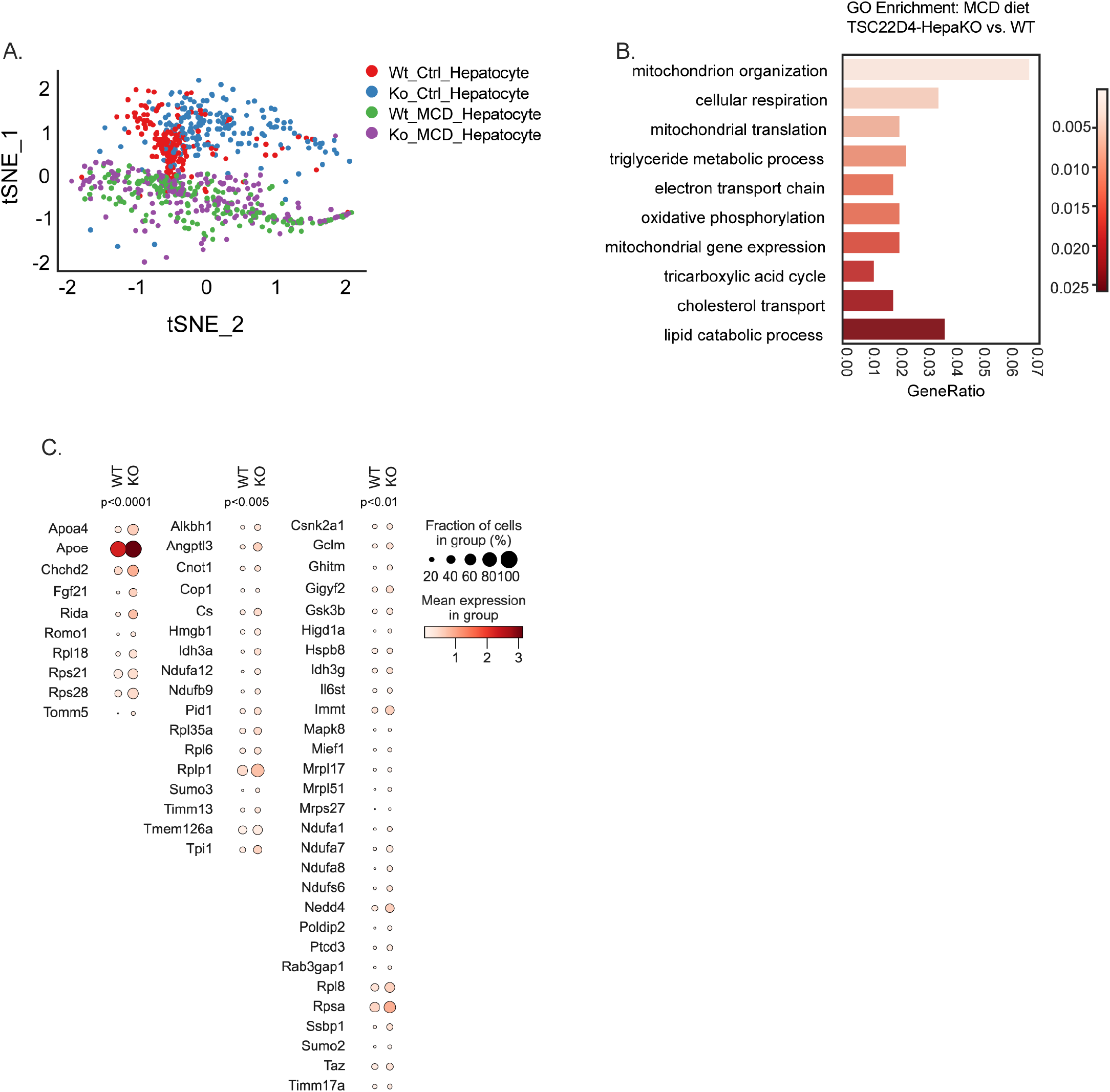
Gene signature in single cell RNAseq from TSC22D4-HepaKO and WT mice. T-distributed stochastic neighborhood embedding (tSNE) analysis of hepatocyte fraction from single cell RNAseq of TSC22D4-HepaKO and WT mice on control or MCD diet (A). Ten of top 100 pathways identified by gene ontology (GO) analysis (B). A dot plot of the genes significantly enriched in TSC22D4-HepaKO cells under MCD diet (C). Abbreviations: MCD, methionine-choline deficient.

Intriguingly TSC22D4-HepaKO mice on MCD diet did not lead to the down-regulation of any genes but rather increased the expression of genes related to mitochondrial complex I and cellular respiration (*Ndufa1,7,8,12, Ndufs6, and Ndufb9*), lipid metabolism (*Apoe, Apoa4* and *Fgf21*) and mitochondrial organization and transport (*Idh3a and g, Timm13* and *Tomm5*). Significantly expressed genes were also measured in a greater percentage of KO cells compared to WT in line with the observation that TSC22D4-HepaKO mice on MCD diet had improved markers of liver damage (Fig. 4C.). Genes contributing to the top 10 pathways identified by GO analysis (Supp. Fig. 4C) were applied to the KEGG pathway database to generate a gene network signature (Fig. 5). KEGG annotated gene clusters identified gene clusters related to NAFLD and oxidative phosphorylation as well as cholesterol metabolism.

**FIG. 5.**
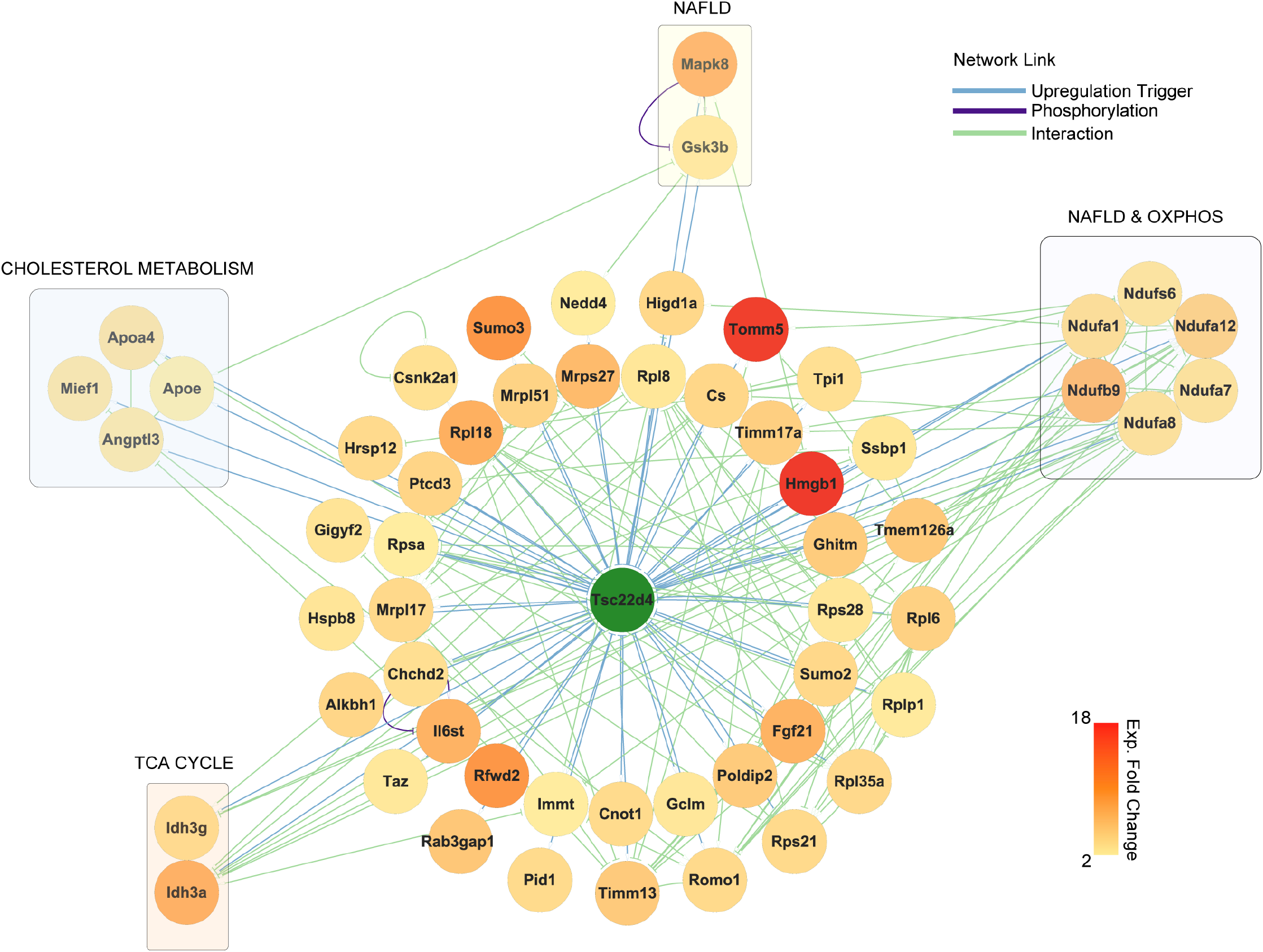
Gene networking reveals mitochondrial maintenance and lipid metabolism. KEGG gene interaction network created from the genes identified in the top ten GO biological pathways.

To evaluate the functional impact of TSC22D4 knock down on mitochondrial respiration in hepatocytes, isolated primary cells were transfected with lipid nanoparticles (LNP) carrying specific TSC22D4-directed siRNA using Arcturus’ LUNAR® delivery technology then analyzed by Seahorse mitochondrial stress tests. In line with our single-cell analysis primary hepatocytes lacking TSC22D4 had increased oxygen consumption rate when compared with siRNA-Ctrl (Fig. 6A) which stemmed from enhanced basal and maximal respiration as well as increased ATP-linked respiration and proton-leak (Fig. 6B). Non-mitochondrial respiration was not significantly altered by TSC22D4 knock down. Together these data support a link between TSC22D4 and mitochondrial function in hepatocytes.

**FIG. 6.**
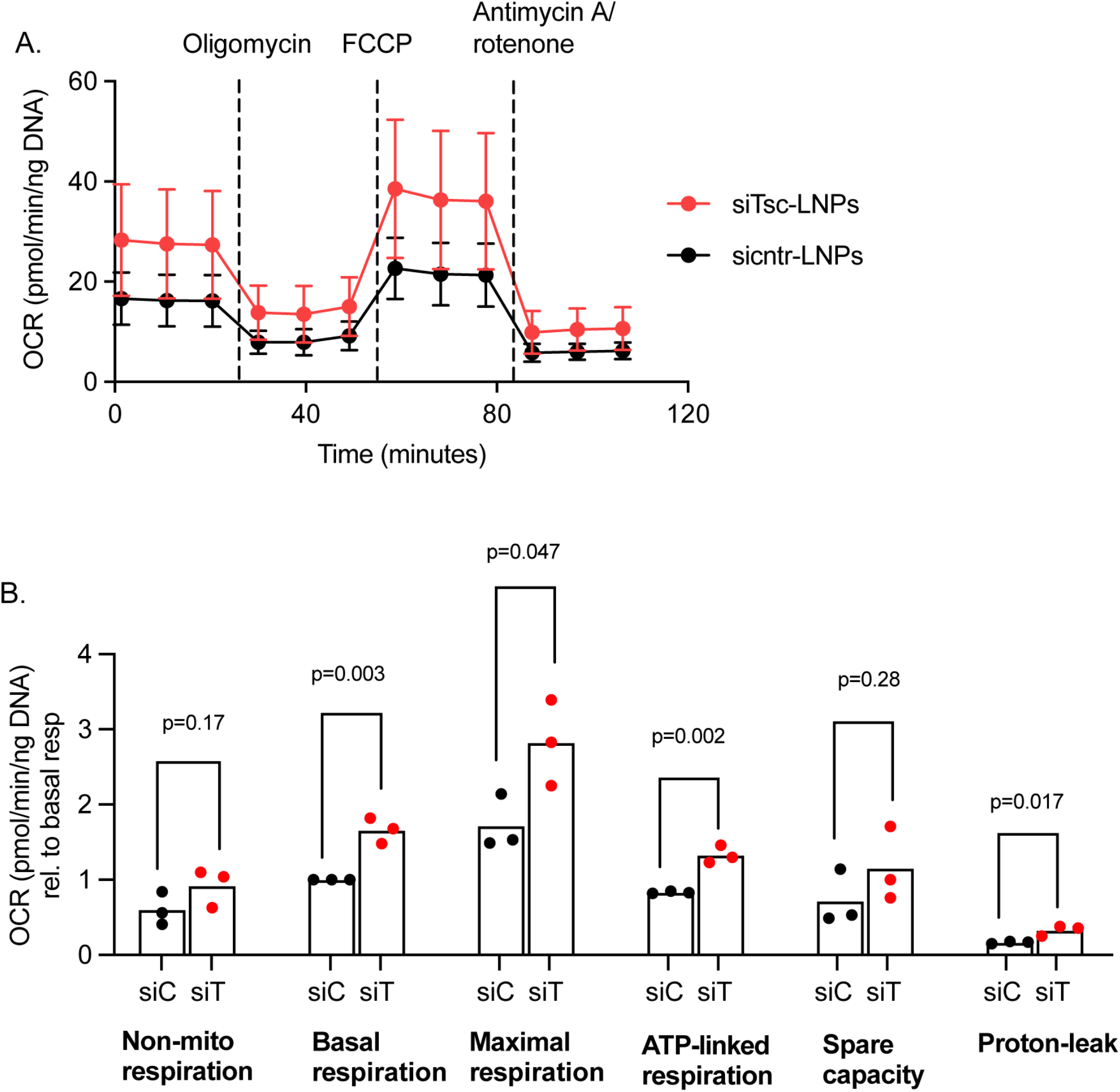
LNP-siRNA mediated knock down of TSC22D4 in primary hepatocytes increases oxygen consumption rate. Seahorse measurements of OCR after 2 days of siControl-LNP or siTSC22D4-LNP knockdown in primary hepatocytes in vitro (n= 10 wells per condition) n=3 independent experiments (A). Metabolic rate calculations from the data in A (B). OCR, oxygen consumption rate

## DISCUSSION

NAFLD is the most common cause of chronic liver disease worldwide [32; 33]. Stages of NAFLD range from early steatosis (NAFL) to non-alcoholic steatohepatitis (NASH) which can lead to irreversible conditions such as cirrhosis or hepatocellular carcinoma. A key event in the progression of NAFLD is lipotoxicity resulting from excessive fat storage in hepatocytes. Distinct biological pathways and molecular mechanisms converge to drive NAFLD progression including oxidative stress, autophagy, apoptosis, and inflammation, ultimately resulting in fibrosis and eventually liver cancer [34; 35].

NAFLD represents the hepatic manifestation of the metabolic syndrome, and indeed the majority of patients affected by obesity-driven type 2 diabetes display signs of liver dysfunction. Of note, NAFLD not only impairs liver function but is also a key driver of systemic insulin resistance as well as macro-vascular complications in long-term diabetes [36], suggesting the existence of hepatic molecular checkpoints to serve as connecting hubs between the intra-hepatic and systemic consequences of liver metabolic dysfunction.In this respect, our previous work has identified hepatic TSC22D4 as a key regulator of systemic insulin sensitivity and hepatic lipogenesis. Hepatocyte-selective knock down of TSC22D4 in type 2 diabetic mice improved glucose metabolism by reducing circulating glucose levels, increasing insulin sensitivity and Akt phosphorylation in the liver [11]. As shown in this study, TSC22D4 expression interestingly also positively correlated with circulating glucose, insulin, and C-peptide levels in patients with steatosis or steatohepatitis (Supp. Fig. 2A-C), further illustrating the importance of regulating TSC22D4 in the context of diabetes-associated liver complications. Also, in a model of cancer-induced wasting, TSC22D4 reduction was sufficient to induce VLDL secretion from the liver and improve tumor-cell mediated metabolic dysfunction in part by modulating lipogenesis in hepatocytes [10].

Our results now support the hypothesis that cell specific targeting of hepatic TSC22D4 may provide a therapeutic strategy to diminish the early adverse events before NASH or other progression occurs, thereby not only ameliorating glucose tolerance and insulin sensitivity but also diminishing the degree of hepatic fibrosis as a long-term consequence of diabetes-related metabolic dysfunction.

Single-cell transcriptomics analysis demonstrated a significant improvement of mitochondrial complex I gene expression, correlating with enhance oxygen consumption rates, upon TSC22D4 loss-of-function. Of note, mitochondrial impairments represent key feature of NAFLD, varying in different stages of the disease [18; 37]. Thus, the hepatocyte-selective activity of TSC22D4 appears to target one of the critical cellular events in the progression of NAFLD.

In addition, some data suggest mitochondrial dysregulation to be an initial step in the pathway to insulin resistance and hepatic steatosis by highlighting the role of mitochondrial complexes in obesity and progression to NAFLD [38-40]. While others have demonstrated deficits in specific complex subunit proteins such as *Ndufa 9* [41] *Ndufa (1, 2, 11), Ndufb (6, 7, 9)* in progressive NAFLD and NASH [42].

Little is known about TSC22D4 or other TSC22-family members in relation to mitochondrial function. One report described four forms of TSC22D4 in developing cerebellar granule neurons (CGN) and identified the 67kDa modified form to be specifically enriched in the mitochondria of differentiated CGNs [43]. TSC22D4 has been shown to heterodimerize with apoptosis inducing factor (AIF), a key protein in mitochondrial redox metabolism [44].

As an outlook, our data suggest that the hepatocyte-selective targeting of TSC22D4 may provide a new path forward to improve systemic insulin sensitivity and glucose handling as well as improving mitochondrial function, thereby preventing and/or reversing even diabetes-related liver fibrosis.

## ABBREVIATIONS

TSC22D4: Transforming growth factor beta-like stimulated clone 22 domain 4
scRNA-seq: single cell RNA-sequencing
TG: triglycerides
MCD: methionine-choline deficient
Met: metformin
LNP: lipid nanoparticle
CCl_4_: carbon tetrachloride

## AUTHOR CONTRIBUTIONS

GW contributed to the investigation, formal analysis, visualization, and writing the original draft. PW and AMh contributed formal analysis, visualization, reviewing and editing the manuscript. AW, MT, IKD, and KY contributed to the investigation. AZ, MR, NV, TH, FDAR, JVR, AF, RM, PL, KT, PK, JP, CPMJ, and JS contributed resources. MS, AMa, and BEU contributed to writing the original draft. TP and LW contributed resources and investigation. PN and JS contributed to the conceptualization, funding acquisition and supervision. SH contributed to the conceptualization, funding acquisition, supervision and writing the original draft.

## ACKNOWLEDGEMENTS

We would like to thank Bettina Walter and the Center for Model System and Comparative Pathology for their technical expertise (human and mouse disease), the IBF and clinical experimental area of Heidelberg University for technical expertise (mouse studies), the Helmholtz Center München (HMGU) Genomics Core facility and Inti Velazquez, Ana Alfaro for scientific discussions and expertise in mouse disease models, Julia Zuber for expertise with Seahorse analysis, and Luke Harrison for editorial contributions.

## SUPPLEMENTAL DATA

**Supplement FIG. 1.**
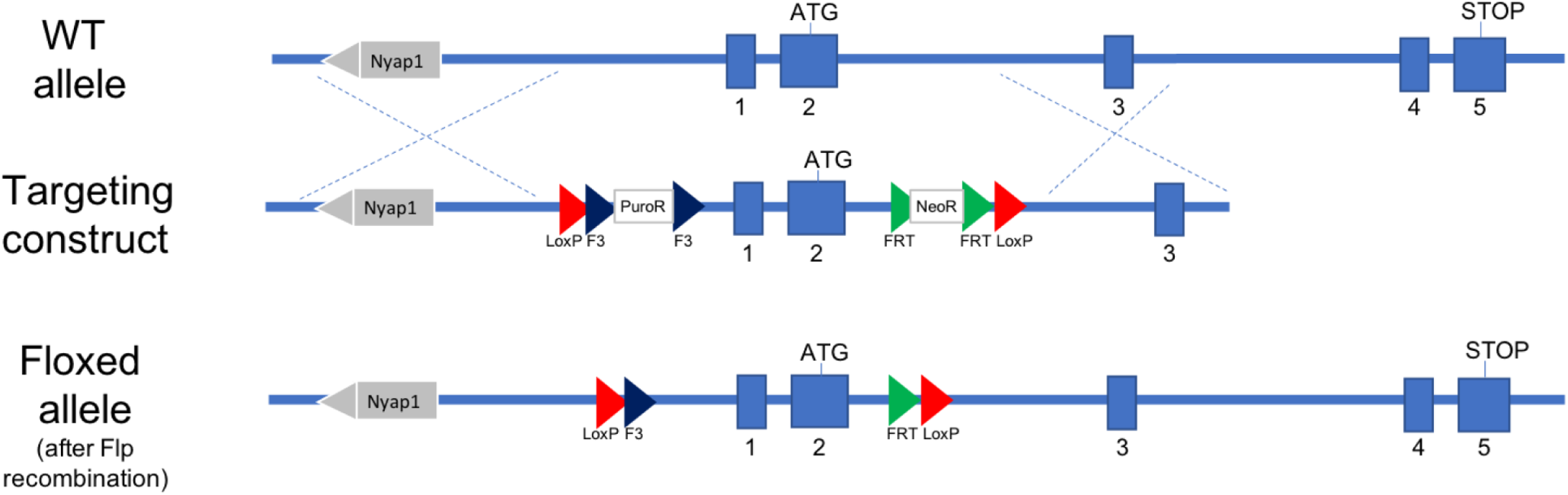
Generation of the TSC22D4 floxed mouse for conditional gene knockout. Graphical representation of the targeting strategy for TSC22D4.

**Supplement FIG. 2.**
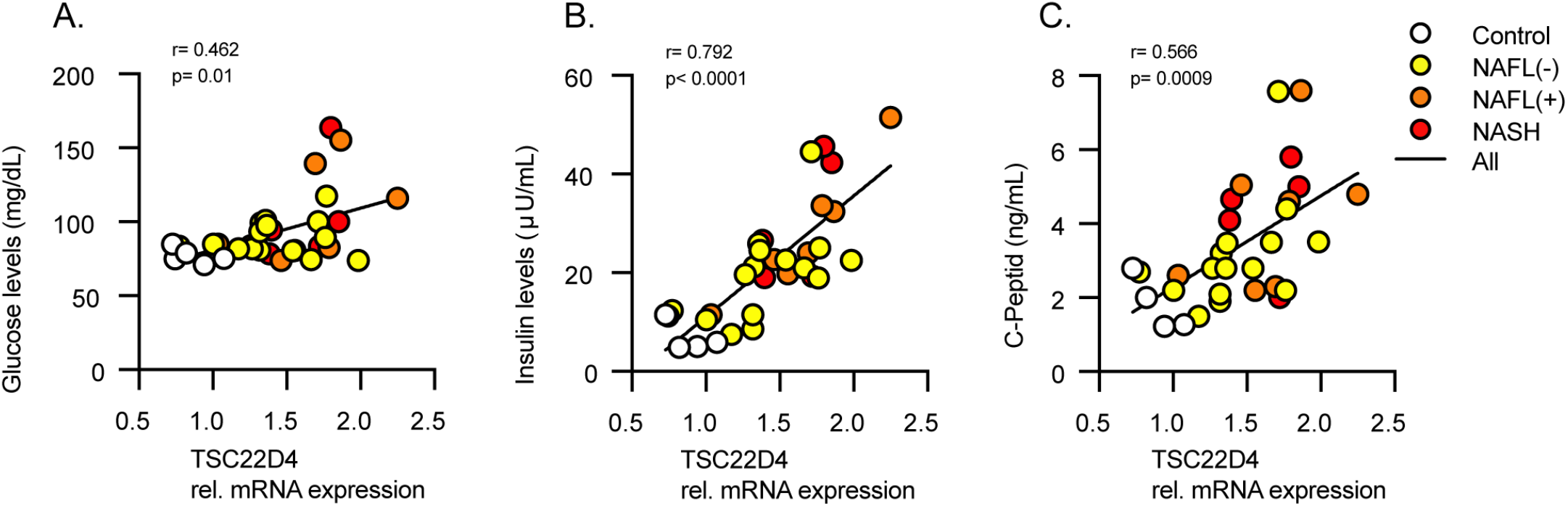
Hepatic TSC22D4 expression correlates with systemic glucose, insulin, and C-peptide levels in patients with steatosis or steatohepatitis. TSC22D4 mRNA expression in liver positively correlated with circulating blood glucose (A), insulin (B) and C-peptide (C) levels.

**Supplement FIG. 3.**
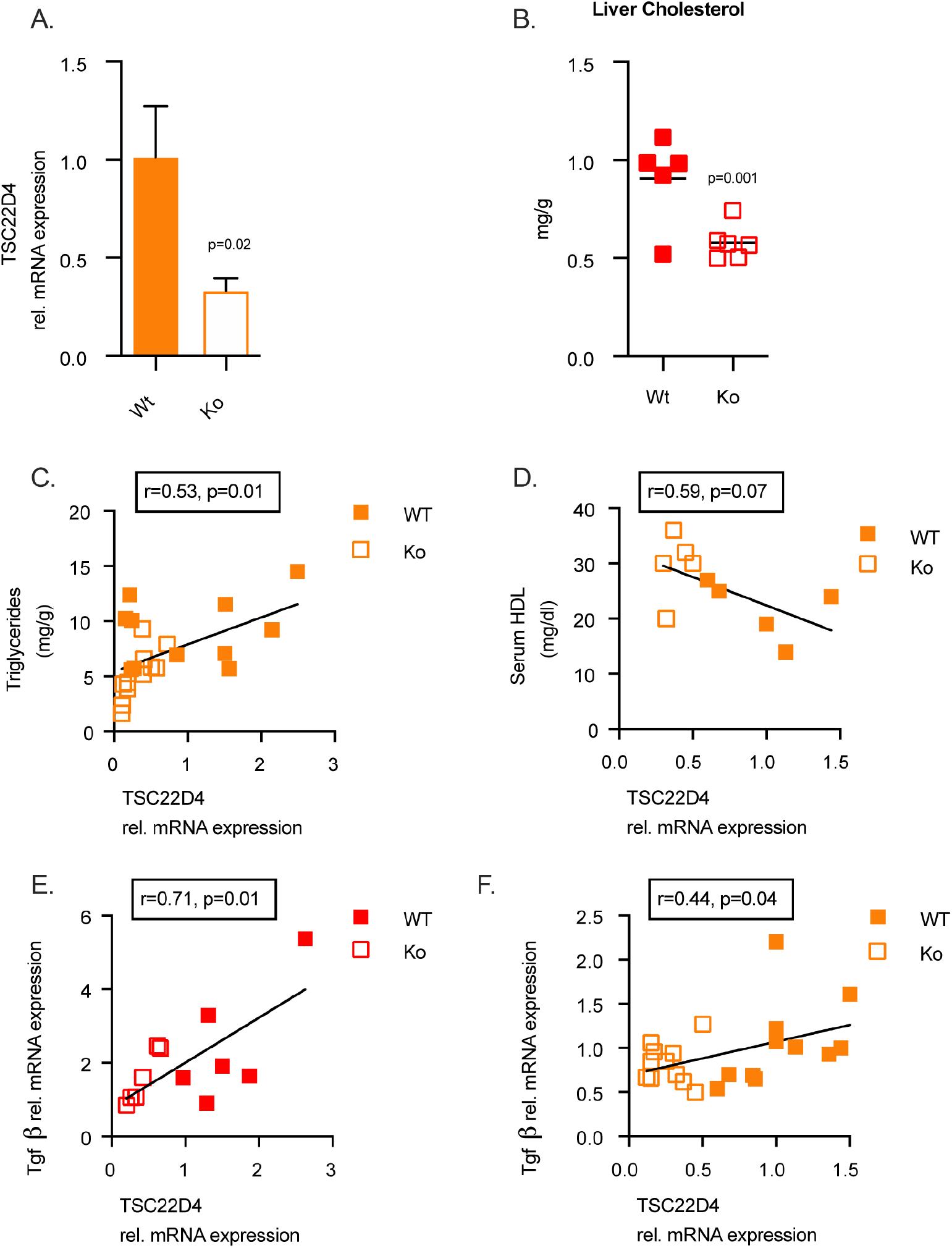
TSC22D4 expression correlates with markers of progressive fatty liver disease in NASH mouse models. TSC22D4 mRNA levels in TSC22D4-HepaKO or WT mice on MCD diet for 3 weeks (A). Hepatic total cholesterol in STAM model (B). TSC22D4 mRNA positively correlated with liver triglycerides (C) and negatively correlated with HDL-cholesterol (D). Hepatic TSC22D4 expression correlated with TGFβ, a marker of NASH progression, in both STAM (E) and MCD diet fed mice (F). Abbreviations: MCD, methionine-choline deficient; CD, control diet; STAM, STAM™ model of low-dose streptozotocin with high fat diet

**Supplement FIG. 4.**
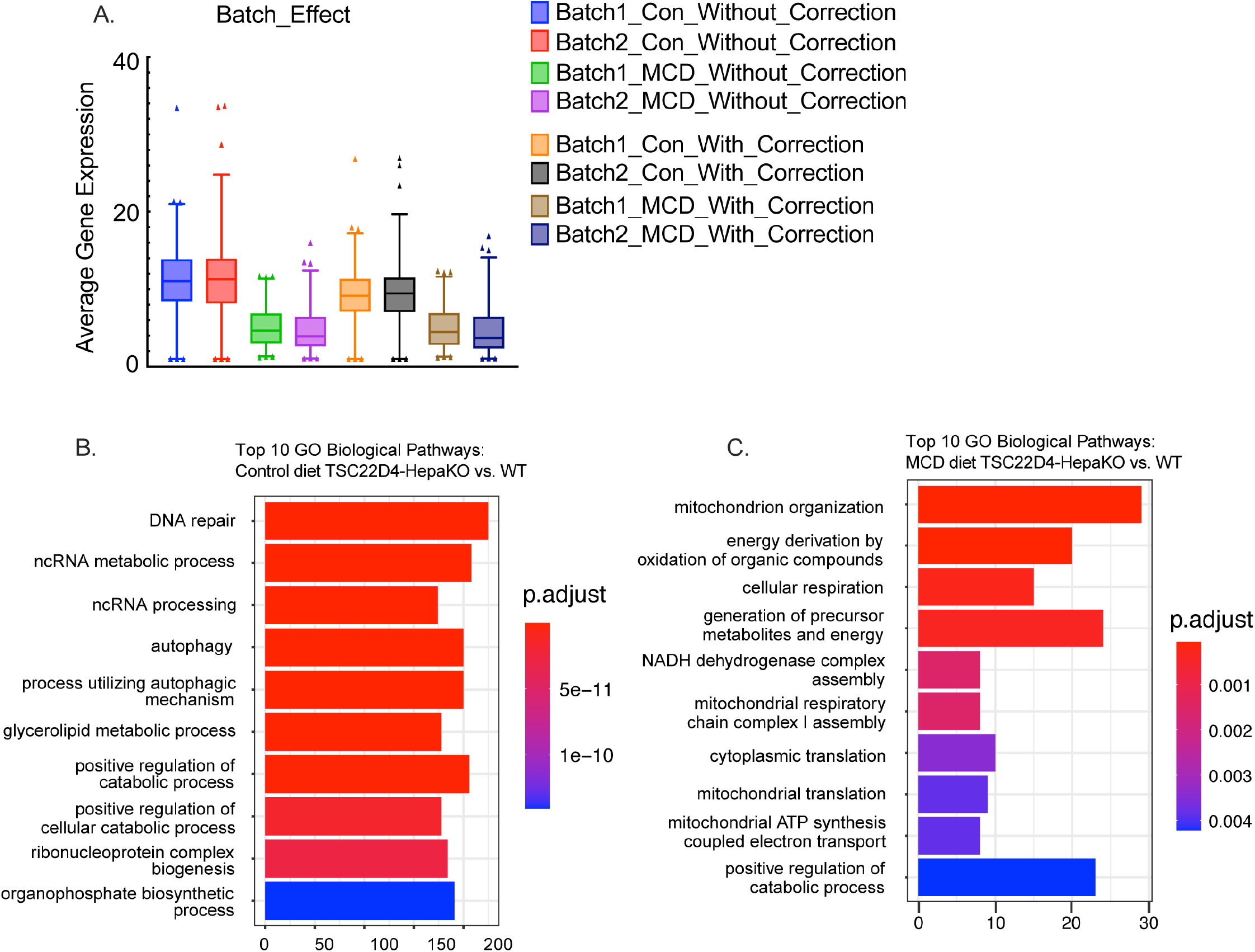
Single cell RNA sequencing of TSC22D4-HepaKO and WT hepatocytes. (A) Batch effect plot (B) Top 10 biological pathways identified by GO analysis in controls. (C) Top 10 biological pathways identified by GO analysis in MCD diet.

